# Whole-genome duplication underlies conserved sexually biased expression of meiotic cohesin genes unique to the teleost fish lineage

**DOI:** 10.64898/2026.06.26.731870

**Authors:** Taiki Niwa, Mariko Kikuchi, Minoru Tanaka

## Abstract

Meiosis is a fundamental process in producing both sperm and eggs, yet recombination landscapes often exhibit sexual differences, known as heterochiasmy. Since meiotic proteins are generally expressed in both sexes, the molecular mechanism driving heterochiasmy remains elusive. The α-kleisin subunit gene of meiotic cohesin, *Rec8*, is expressed bisexually in mammals, while its putative teleost ortholog, *rec8a,* is expressed in a female-biased manner, presumably due to the presence of its paralog originating from the teleost-specific whole-genome duplication (TGD). Here, we elucidated the evolutionary history and expression dynamics of α-kleisin genes across teleost lineages. Through comprehensive phylogenetic and synteny analyses, we revealed that major teleost lineages retain two copies of *rec8* and *rad21*, with *rec8* loci experiencing drastic chromosomal rearrangements immediately after the TGD. Using *in situ* hybridization and single-cell transcriptome data in medaka and zebrafish, we demonstrated a conserved sexually biased expression pattern: *rec8a* is predominantly female-biased, whereas *rec8b* exhibits male-biased expression during gametogenesis. Furthermore, comparative epigenetic analyses revealed that the conserved sexually biased expression is driven by lineage-specific *cis*-regulatory elements, rather than conserved ones. Motif analyses imply that regulatory rewiring by transcription factors, including *foxl2l* in particular, might have played a crucial role in the establishment and maintenance of this paralog divergence. Our findings highlight how whole-genome duplication and subsequent genomic and epigenetic rewiring subdivided the bisexual function of *rec8*, offering insights into sexually distinct meiotic regulation.

**Highlights:**

- Teleosts possess a unique α-kleisin repertoire originating from the TGD.
- Teleost *rec8* paralogs exhibit conserved sex-biased expression during meiosis.
- Drastic genomic rearrangements after the duplication rewired the teleost *rec8* loci.
- The conserved expression pattern is governed by lineage-specific CREs.
- Those CREs harbor similar types of TFBSs such as Fox-family TFs.

**Graphical abstract:** 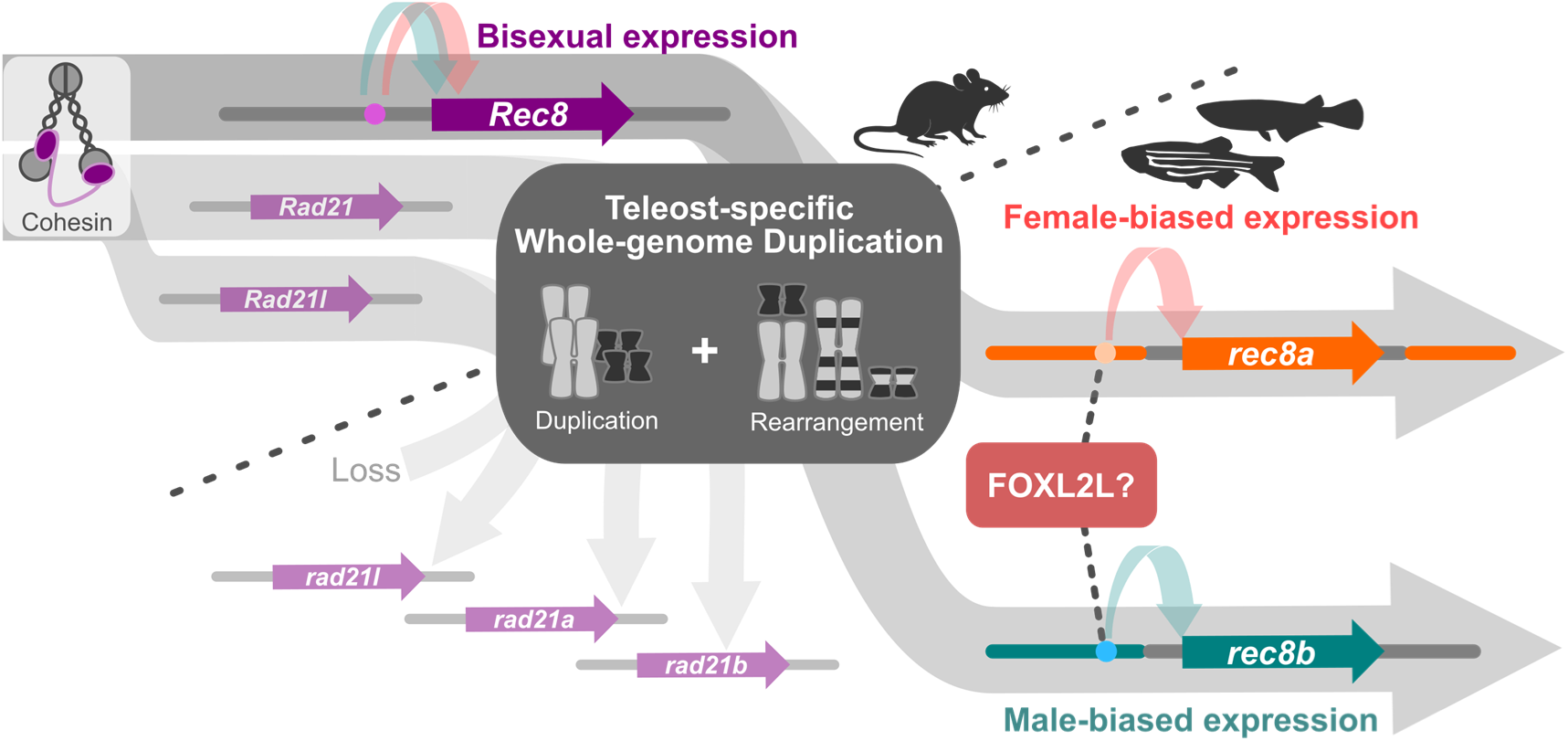

## Introduction

Meiosis is a common process to produce both eggs and sperm. Although the consequent gametes exhibit huge morphological differences, both eggs and sperm undergo homologous recombination and two successive cell divisions in the same manner. Thus, major proteins acting in meiosis are essentially expressed and function in both sexes. When homologous chromosomes recombine, these chromosomes are cleaved, scaffolded, synapsed, and repaired, and all these processes are indeed mediated by various types of proteins dedicated to each process and are not sex-specific (Zickler and Kleckner, 2023).

At the same time, previous studies have uncovered robust sexual differences in recombination landscapes, namely the rate and localization of crossovers, which is termed heterochiasmy (Morgan, 1914; Kondo et al., 2001; Kochakpour and Moens, 2008; Giraut et al., 2011; Sardell et al., 2018; Cahoon and Libuda, 2019). This phenomenon is widely observed in sexually reproducing organisms including both animals and plants, regardless of monoecy and dioecy, showcasing its fundamental property of meiosis under sexual reproduction (Cahoon and Libuda, 2019). Corresponding to heterochiasmy, the lengths of chromosomal axes, proteinaceous structures lining between paired homologous chromosomes, also vary between sexes (Kochakpour and Moens, 2008; Gruhn et al., 2013; Zickler and Kleckner, 2023). These lengths are recognized to reflect the differences in meiotic chromosomal structure, namely loop sizes in the loop-axis arrangements of meiotic chromosomes, and are associated with the number of crossovers (Gruhn et al., 2013; Wang et al., 2017). Different chromosomal structures potentially have influence on the accessibility of the recombination machinery, further implying its causal relationship with heterochiasmy (Gruhn et al., 2013; Song et al., 2021; Capilla-Pérez et al., 2021). Despite these accumulating study cases, the underlying molecular mechanism of these phenomena remains elusive.

We previously identified the *rec8a* gene in Japanese medaka (*Oryzias latipes*) as a potential clue for this unsolved mystery. *rec8a* is identified as one of the direct target genes of a transcription factor, *foxl2l* (formerly termed as *foxl3*), which regulates the germline sexual identity of medaka through its female-biased expression (Nishimura et al., 2015; Kikuchi et al., 2020). Owing to this gene regulatory relationship, *rec8a* also exhibited female-specific expression in developing gonads and its null mutation induced by genome editing resulted in female-specific sterility (Kikuchi et al., 2019, 2020). Medaka *rec8a* is predicted as an ortholog of mammalian *Rec8*, which encodes an α-kleisin subunit of a cohesin complex. These complexes play crucial roles in meiosis by scaffolding the loop-axis structures of chromosome axes for homologous recombination and by bundling sister chromatids to ensure the proper timing of meiotic chromosome segregation (Ishiguro, 2019). Mammals have three α-kleisin genes: *Rad21*, *Rad21l*, and *Rec8*, each of which exhibits distinct expression pattern as represented by the meiosis-specific expression of *Rad21l* and *Rec8* (Ishiguro et al., 2014; Ishiguro, 2019). Notably, these mammalian genes are expressed in both sexes in contrast to medaka *rec8a*. In addition, other genes involved in cohesin complexes including the putative *rad21*, *rec8*, *sgol1,* and *nipbl* orthologs were also upregulated or downregulated in medaka ovaries after *foxl2l* knockout (Kikuchi et al., 2019). These observations shed light on the significance of sexually biased cohesin expression during gametogenesis in medaka.

Interestingly, several teleost species including zebrafish (*Danio rerio*) and Nile tilapia (*Oreochromis niloticus*) possess duplicated copies of *rec8* as briefly mentioned in previous reports (Crespo et al., 2019; Luo et al., 2021; Imai et al., 2021; Hsu et al., 2025). Furthermore, these *rec8* paralogs in the Nile tilapia had sexually biased expression as seen in medaka *rec8a* (Luo et al., 2021). This specific situation, found only in teleosts, is presumably attributed to the whole-genome duplication experienced in the teleost common ancestor about 300 million years ago (TGD), which is considered to have shaped unique gene repertoires and functions in teleost lineages (Hoegg et al., 2004; Braasch et al., 2016; Pasquier et al., 2017). This duplication might have led to a relaxed evolutionary constraint of the bisexual expression of the ancestral *rec8* paralogs. However, this assumption is still equivocal due to the limited genomic information used in the previous study, hindering a comprehensive view of the teleost α-kleisin repertoire and their actual origin.

In this study, we aimed to profile the gonadal expression pattern of teleost α-kleisin genes and elucidate the root of the unique expression pattern in teleost *rec8* paralogs. Our analyses involving molecular phylogeny, gene expression, and epigenetics uncovered the complete repertoire of teleost α-kleisin genes, the conserved sexually biased expression of two *rec8* paralogs, and its potential origin in the TGD and its subsequent chromosomal rearrangements.

## Results

### Molecular phylogeny of jawed vertebrate α-kleisin genes

To resolve the ambiguous α-kleisin phylogeny in vertebrates, we compared both amino acid sequences and synteny in the genomic regions surrounding α-kleisin genes. In homolog searches, two copies of the *rec8* and *rad21* genes were identified in a wide range of teleost genomes. Each of these four homologs, henceforth *rec8a, rec8b, rad21a,* and *rad21b,* formed a monophyletic clade on their molecular phylogenetic trees, except for the *rad21* orthologs in three species (Asian bonytongue, *Paramormyrops kingsleyae*, and European conger) (Fig. 1a, b). In contrast, *rad21l* was always detected as a single copy in all the vertebrate genomes even in teleosts. In the phylogenetic tree including both *rad21* and *rad21l, rad21l* formed its own clade branching from *rad21* after the divergence of lamprey *Rad21* orthologs but before the divergence of cartilaginous fish *Rad21* orthologs (Fig.1c). We then examined the gene arrangements on the genomic regions around these α-kleisin genes. Synteny analyses revealed similar arrangements of paralogs in two different loci of teleost genomes: *rad21a* / *rad21b* loci and *snpha* (with *rad21l*) / *snphb* loci (Fig. 1d). Genomic regions corresponding to these *rad21a/b* and *rad21l* loci were identified as single loci in tetrapod genomes (Fig. 1d). On the other hand, *rec8* homologs did not show this pattern, where genes flanking *rec8a* and *rec8b* were not paralogous (Fig. 1e). The teleost *rec8b* and tetrapod *Rec8* loci partly shared their flanking genes including *ipo4* and *tm9sf1*, while the teleost *rec8a* and tetrapod *Rec8* loci did not share any flanking genes (Fig. 1e). The teleost *rec8a* locus had a paralogous counterpart in another locus of teleost genomes, where a paralogous arrangement of *rec8a* flanking genes was detected with the exception of *rec8a* itself (Sup. Fig. 1). These loci were further orthologous to a single locus in tetrapod genomes, while *rec8* was absent there (Fig. 1e). Medaka *rad21b*, Asian bonytongue *rec8b*, and *rec8b* of Siluriformes species (catfishes) were absent from these genomes even when considering these syntenic correspondences.

**Figure 1.**
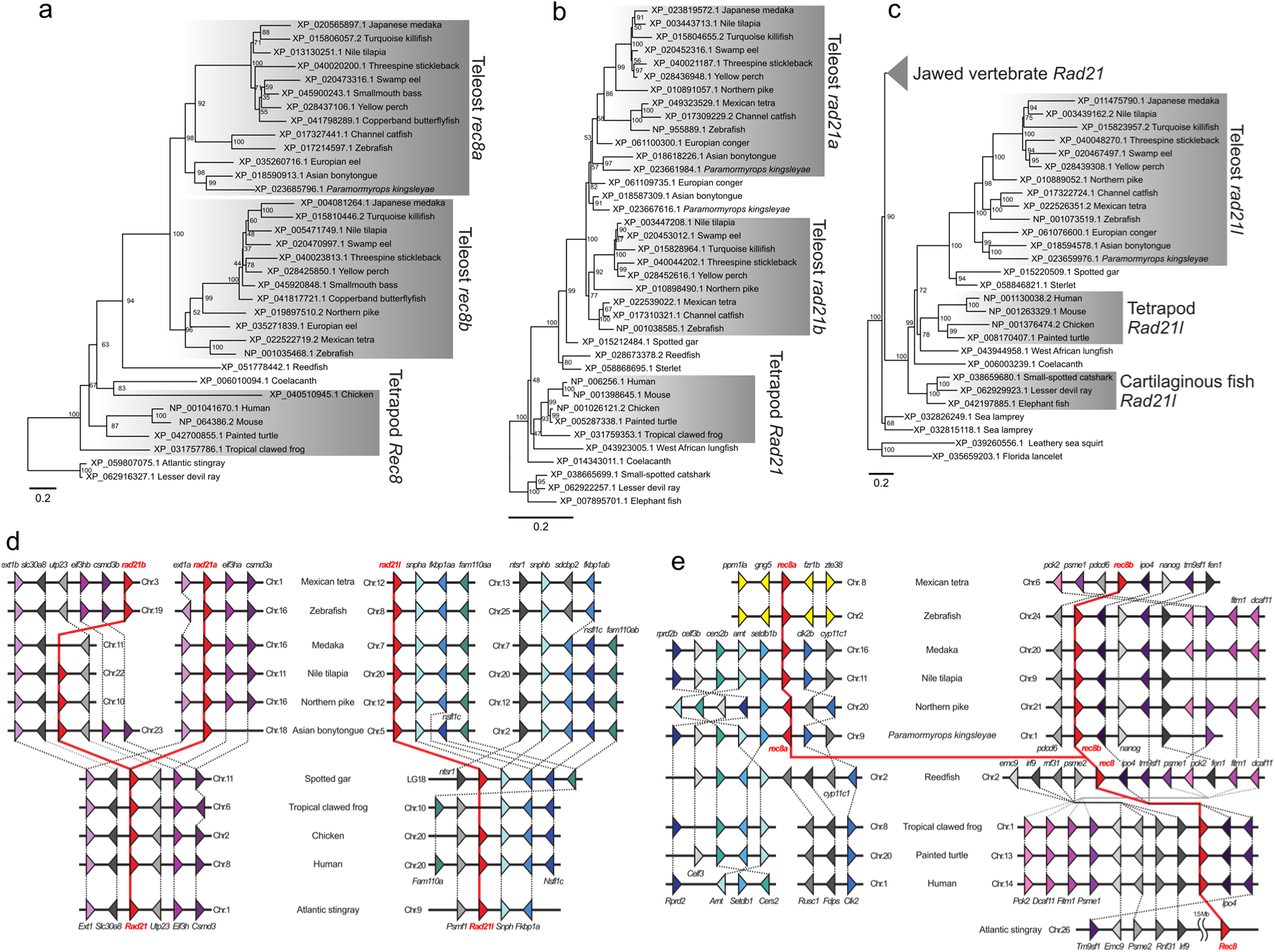
Molecular phylogeny and conserved synteny of vertebrate α-kleisin genes. **(a-c)** Phylogenetic trees of jawed-vertebrate *rec8* (a), *rad21* (b), and chordate *rad21-rad21l* (c) homologs built by the maximum likelihood method. Values on nodes indicate the results of 1000-times bootstrap tests. (**d, e)** Schematic diagrams showing colinear syntenic arrangements of α-kleisin gene loci. Genes and their transcriptional directions are represented as triangles, and their homology assumed from common functional gene annotations was depicted as edges. Adjacency of genes in this diagram shows their relative positions on the genomes, rather than the actual side-by-side arrangements.

### Expression patterns of teleost α-kleisin genes in male and female germ cells

Based on this gene repertoire, we examined localizations of medaka *rec8a*, *rec8b*, *rad21a*, and *rad21l* transcripts using *in situ* hybridization on testicular and ovarian tissue sections. These gonadal tissues contained germ cells in the early stages of gametogenesis, including both mitotic and early meiotic germ cells, distinguished by nuclear morphology (e.g., the presence of prominent nucleoli in the center of mitotic germ cell nuclei). *rec8a* was expressed only in a subset of mitotic germ cells in both males and females with more prominent expression in females than in males (Fig. 2). In contrast, *rec8b* was expressed in a subset of mitotic germ cells solely in males, but completely lacked expression in females (Fig. 2). *rad21a* and *rad21l* were expressed in mitotic germ cells and a subset of meiotic germ cells in males, whereas these were detected in most female germ cells including developing oocytes (Fig. 2).

**Figure 2.**
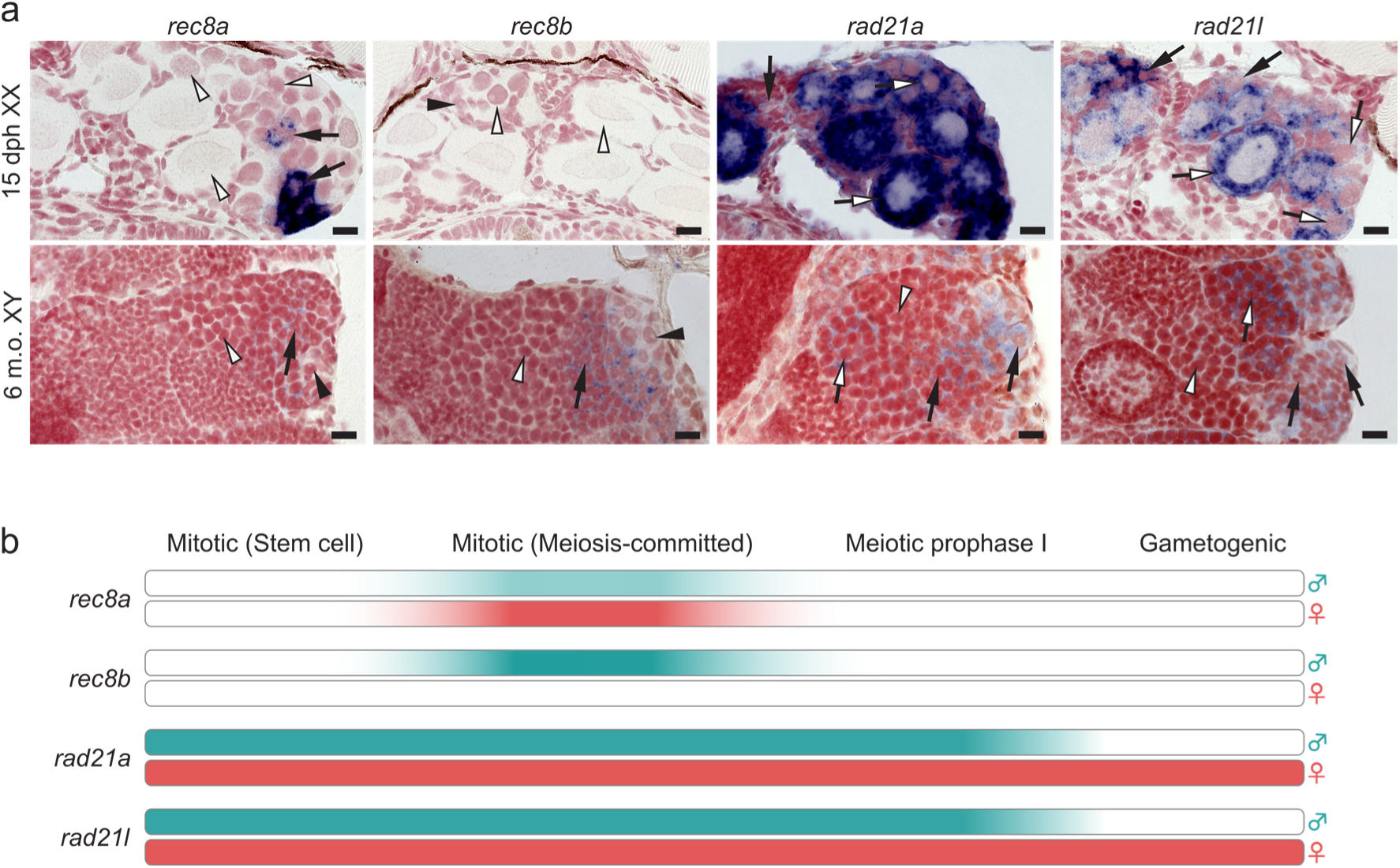
Spatial expression patterns of α-kleisin genes in medaka gonads. **(a)** Localizations of α-kleisin transcripts on cross sections of adult (6 m.o.) testes and 15 dph developing ovaries in medaka. 3–5 individuals were used for each condition; results were consistent among replicates. Probes were detected as purple insoluble precipitates and nuclei were detected as red background staining on the tissue sections. Arrows and arrowheads show cells with and without signals, respectively. The black and white color of an arrow / arrowheads shows different cell types; mitotic and meiotic germ cells, respectively. Scale bar: 10 μm **(b)** Schematic diagram showing temporal expression patterns of α-kleisin genes in medaka germ cells.

To further assess conservation of the expression patterns among teleosts, we utilized two previously reported single cell RNA-seq (scRNA-seq) data derived from a zebrafish testis and ovary (Liu et al., 2022; Qian et al., 2022). The clustering analysis based on transcriptomic profiles successfully identified distinct cell types, as represented by the expression of cell-type-specific markers: *nanos2*, *dmc1*, *zp3b* in females and *ddx4*, *dmc1*, *odf3b* in males, which were also used in the original studies (Fig. 3a, b). Notably, *nanos2* or *ddx4*-positive cells were annotated as mitotic germ cells, and *dmc1*-positive cells were annotated as meiotic germ cells (prophase I). *rec8a* was expressed only in a subset of female mitotic germ cells (Fig. 3c, d). This expression was limited to a later stage of mitotic germ cells which are recognized as meiosis-committed mitotic germ cells (labelled as pre-meiotic in Fig. 3). *rec8b* was expressed from mitotic to meiotic phases in both male and female germ cells, yet its peak expression level and the proportion of *rec8b*-expressing cells was clearly higher in males (Fig. 3c, d). Although zebrafish possessed both *rad21a* and *rad21b*, only *rad21a* was expressed in the zebrafish gonads (Fig. 3c, d). *rad21a* expression was observed in most cell types analyzed here, but substantially more intense in mitotic germ cells in both sexes and early oocytes (Fig. 3c, d). *rad21l* was expressed in both mitotic and meiotic germ cells with its peak expression level at meiosis (Fig. 3c, d). *rad21l* expression in females continued until early oocytes, similar to *rad21a*.

**Figure 3.**
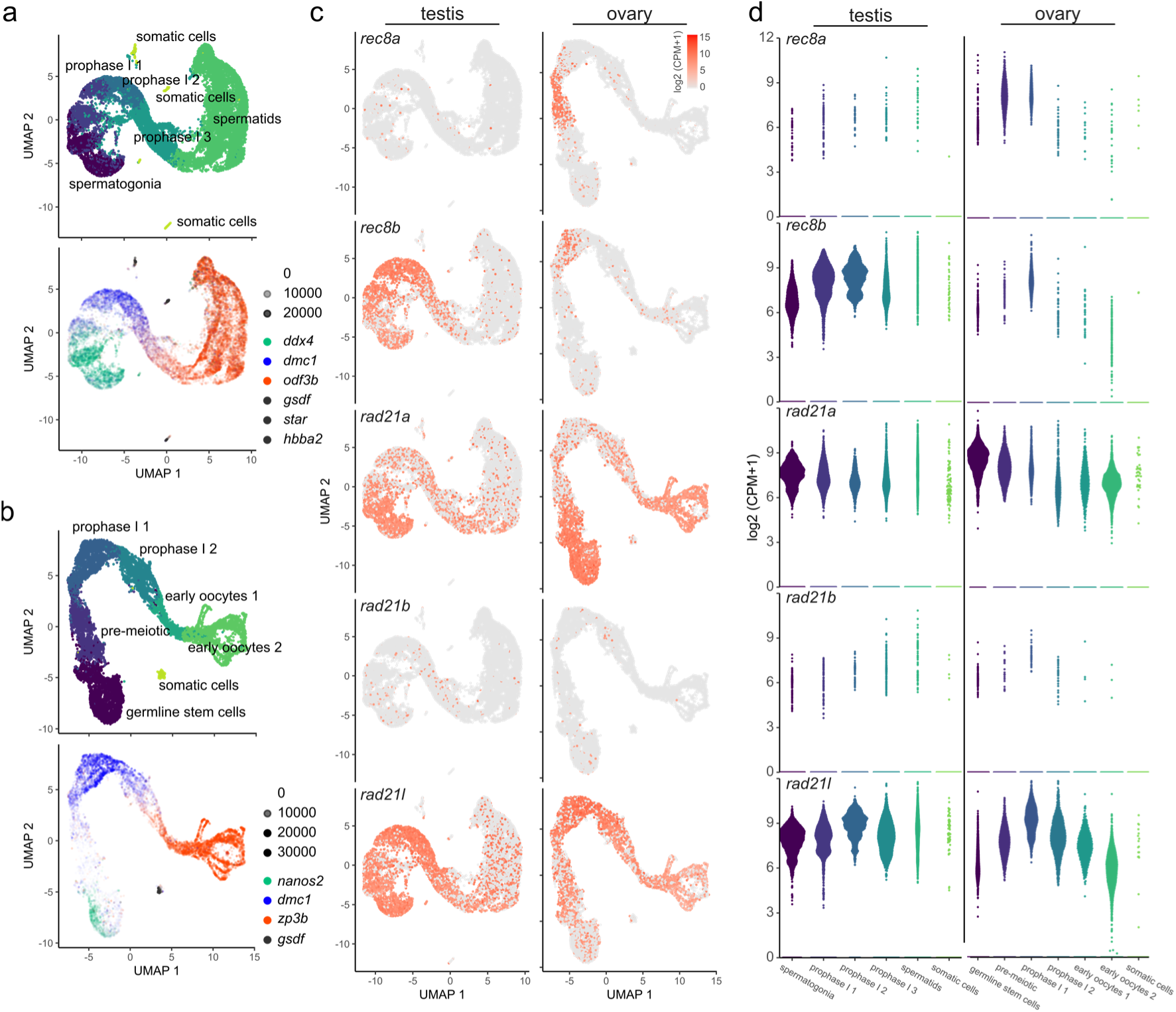
Expression patterns of α-kleisin genes in zebrafish gametogenesis. (a, **b)** Visualization of transcriptomic profiles derived from zebrafish testicular (a) and ovarian (b) scRNA-seq in UMAP (uniform manifold approximation and projection) plots. Cell types were manually annotated based on the clustering results by cell-type-specific markers as represented by the *ddx4* (spermatogonia), *nanos2* (germline stem cells), *dmc1* (meiotic prophase I), *odf3b* (spermatids), *zp3b* (oocytes), *gsdf*, *star*, *hbba2* (somatic cells), and coded by different colors. Expression levels of marker genes were represented by counts per million (CPM) in a cell. **(c, d)** Expression levels of α-kleisin genes on UMAP plots (c) and in each cell type (d). Expression levels were transformed to log2 (CPM + 1).

### Putative regulatory mechanisms of mouse *Rec8* and zebrafish *rec8a/b*

Considering the discrepancy in the synteny around the homologous *Rec8* loci between teleost and non-teleost vertebrates (Fig. 1e), these differences might have influenced their expression patterns. Teleost *rec8b* and tetrapod *Rec8* did not share the upstream adjacent gene; *rec8b* is adjacent to *pdcd6* while tetrapod *Rec8* is adjacent to *Irf9*, which implies an ancient chromosomal rearrangement occurred at this intergenic region. This rearrangement potentially resulted in differential transcriptional regulation of *rec8* paralogs through *cis*-regulatory mechanisms. To assess the gene-regulatory potential of these regions, we identified putative *cis*-regulatory elements (CREs; e.g. enhancers and promoters) of these regions by using public datasets of ATAC-seq (assay for transposase-accessible chromatin using sequencing) and ChIP-seq (chromatin immuno-precipitation followed by sequencing) for active histone marks (H3K27ac and H3K4me3) derived from mature testes of both mouse and zebrafish (Menon et al., 2019; Spruce et al., 2020; Yang et al., 2020). Additionally, sequence conservation around putative CREs was assessed by overlaying phyloP scores derived from the whole-genome alignment of 241 mammalian species or the regional genome alignment of 43 actinopterygian fish species.

The mouse *Rec8* upstream region included one major and two minor ATAC-seq peaks flanked by both H3K27ac and H3K4me3 marks. This suggests that these sites are accessible during spermatogenesis and potentially serve as CREs of the downstream *Rec8* gene (Fig. 4a). These putative CREs were further validated by their overlaps with the ChIP-seq peaks of retinoic acid receptors (RARs) and DMRT1 with their corresponding binding motifs, both of which were well characterized as transcription factors involved in meiotic initiation in mouse (Sup. Fig. 2) (Bowles et al., 2006; Matson et al., 2010; Koubova et al., 2014). Zebrafish *rec8a* and *rec8b* loci harbored one and two putative CREs in their upstream, respectively (Fig. 4b, c). The most accessible sites among these putative CREs were those around the transcription start sites (TSS) of the *rec8* homologs. Along with this, these major CREs at the TSSs were relatively conserved compared with surrounding non-coding regions and with the minor CREs, according to the phyloP scores (Fig. 4 and Sup. Figs. 3 and 4). On the other hand, these major CREs in zebrafish were only alignable with species within Cypriniformes and not with the other teleost lineages (Sup. Fig. 3). Similarly, short stretches within the 5’UTRs of medaka *rec8a* and *rec8b* exhibited relatively high conservation scores, while they were only alignable with a subset of teleost lineages (Sup. Fig. 4).

**Figure 4.**
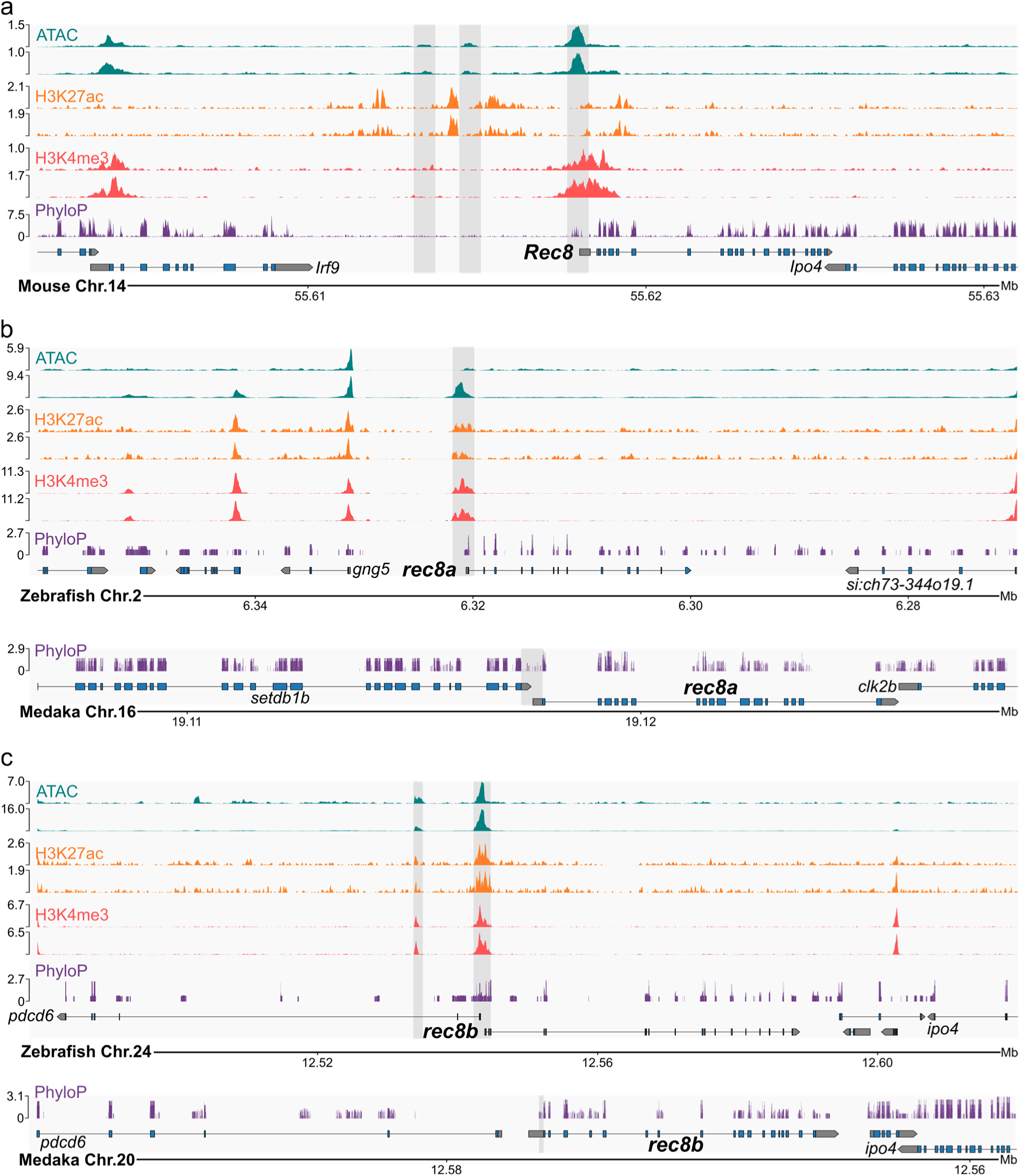
Discrepancy in *cis-*regulatory elements of *Rec8* homologs. **(a-c)** Profiles of chromatin accessibility (ATAC), two active histone marks (H3K27ac and H3K4me3) in testicular cells, and sequence conservation (PhyloP scores), at the mouse *Rec8* (a), zebrafish and medaka *rec8a* (b), and zebrafish and medaka *rec8b* (c) loci. Two rows of each sequencing result indicate different biological replicates in the original studies. Shadings in the plots show the location of the putative CREs defined mainly by ATAC-seq peaks (zebrafish and mouse) or conserved non-coding regions (medaka). Read counts were normalized as CPM. PhyloP scores for mouse were adapted from the previous calculation for 241 mammalian species by the Zoonomia consortium, and those for zebrafish and medaka were calculated from the regional genome alignments of 43 fish species (see also Sup. Figs. 3 and 4).

Within these putative CREs, we further assessed binding potentials of transcription factors (TFs) by incorporating both position weight matrices of TF motifs and chromatin accessibility as the prior probability of TF-motif presence, and assessed their sequence conservation by phyloP scores. In the mouse *Rec8* locus, RARs were the most likely TF binding to the major CRE consistent with their highest conservation scores, which was in line with the ChIP-seq result above (Fig. 5 and Sup. Fig. 2). Androgen receptors harbor their binding motif at the mouse minor CRE, with a prediction score similar to those of RARs, while their conservation score were low (Fig. 5).

**Figure 5.**
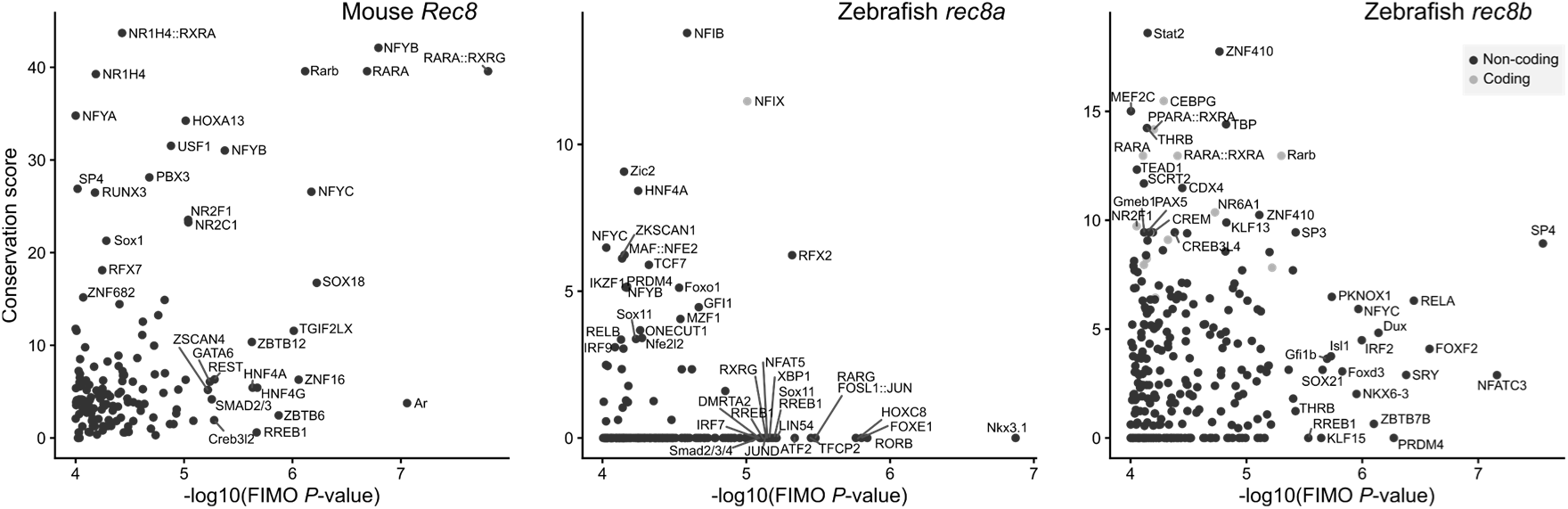
Transcription factor binding sites inferred within the CREs. Horizontal axes show results of TF-binding-site prediction by FIMO with ATAC-seq coverage as its prior distribution. Conservation score represents the sum of phyloP score at a given motif span. TF names were displayed only for those with the top 20 prediction scores or top 20 conservation scores. Dots with pale colors indicate motifs predicted within coding regions, which implies overrepresentation in their conservation scores.

In the zebrafish *rec8a* and *rec8b* CREs, other types of TFs were prominent in the prediction results, when compared with the mouse *Rec8* locus. Although RAR binding sites were predicted at these loci, their prediction scores and/or conservation scores were not as high as those at the mouse *Rec8* locus (Fig. 5). The repertoires and significance of predicted TF motifs were largely different between zebrafish *rec8a* and *rec8b*, while these CREs shared several common TFBSs including those of the forkhead-box (Fox) protein family, for instance (Fig. 5).

## Discussion

### The evolutionary history of teleost α-kleisin genes

Several previous studies reported the presence of duplicated copies of *rec8* and *rad21* in zebrafish and Nile tilapia (Crespo et al., 2019; Luo et al., 2021; Imai et al., 2021; Hsu et al., 2025). However, the origin of the duplication was unclear. With finer taxonomic sampling, we resolved the molecular phylogeny of jawed-vertebrate α-kleisin genes and revealed that a wide range of teleosts possessed five species of α-kleisin genes, namely *rec8a*, *rec8b*, *rad21a*, *rad21b*, and *rad21l* (Fig. 1). Our analyses first validated the orthology of these genes to *rec8*, *rad21*, and *rad21l* in non-teleost vertebrates. Considering the phylogenetic tree topology, emergence of both teleost-specific *rec8* and *rad21* paralogs is reasonably explained by simultaneous duplication events in the teleost common ancestor (Fig. 1a, b). In addition, all the teleost *rad21a* and *rad21b* loci analyzed here exhibited conserved synteny among them, indicating their origin from a regional duplication as a block, rather than local duplication involving only *rad21* gene (Fig. 1d). Considering synchrony of two duplication events and this conserved paralogous synteny, these duplication events most likely correspond to the TGD.

*Rad21l* was first identified as a vertebrate specific paralog for *Rad21* (Gutiérrez-Caballero et al., 2011), yet the precise timing of its emergence remained elusive. Our phylogenetic analysis pinpointed the timing of the *Rad21-Rad21l* divergence after the cyclostome branch and before the cartilaginous fish branch, implying its origin in one of the two consecutive whole-genome duplications in the jawed vertebrate common ancestor (Fig. 1c). Teleost *rad21l* was identified as single copy orthologs unlike *rec8* and *rad21* (Fig. 1c, d). We identified the paralogous counterpart of the *rad21l* locus that lacked the *rad21l* itself in teleost genomes, indicating that the TGD indeed duplicated this locus, yet one of those *rad21l* copies was lost early after the duplication, leading to the single-copy presence of teleost *rad21l*.

In contrast to *rad21* paralogs, *rec8a* and *rec8b* loci did not share any flanking genes. The *rec8a* locus (*setdb1b* locus) had its paralogous counterpart (*setdb1a* locus), while this counterpart was distinct from the *rec8b* locus (Sup. Fig. 1). This complex arrangement indicates that a small fragment of the original *rec8a* locus was translocated into the current locus immediately after its duplication. This arrangement was mostly conserved among teleost lineages; however, *rec8a* of Ostariophysi, including zebrafish, Mexican tetra, and channel catfish, were located at another locus (Fig. 1e). These observations of frequent *rec8a* translocations may indicate the translocation-prone property of this locus, which resembles jumping sex determining loci of *Takifugu* and salmonid fishes (Faber-Hammond et al., 2015; Bertho et al., 2021; Kabir et al., 2022).

Remarkably, most of the drastic chromosomal rearrangements we identified here occurred intensively right after the TGD, and before the emergence of the Eloposteoglossocephala lineage, the first branch from the major teleost lineage among extant teleosts (Parey et al., 2023). Inoue et al. proposed a two-phased evolutionary scenario that both the gene repertoire and genomic arrangements intensively reshaped immediately after the TGD and reached the steady state later on (Inoue et al., 2015). Our observations exactly align with this model, highlighting the magnitude of the TGD with its subsequent effect on the evolutionary traces of teleost α-kleisin genes and their surrounding genomic contexts.

### Unique expression patterns of α-kleisin genes in teleost meiosis

In mouse, three α-kleisin genes are sequentially expressed in different substages of meiotic prophase I (Ishiguro, 2019). RAD21, expressed mostly in mitosis, first decreases its expression in contrast to the upregulation of RAD21L and REC8 around the pre-meiotic DNA replication. RAD21 appears again at a later stage of meiotic prophase I as RAD21L disappears.

Medaka and zebrafish *rad21l* were continuously expressed in a broader span of the gametogenic process than *rec8* paralogs, ranging from the mitotic phase to the later stage of meiotic prophase I (Figs. 2 and 3). In zebrafish, the *rad21l* null mutation caused a male-biased sex ratio; all the male mutants were completely fertile, while female mutants generated by the additional *tp53* mutation were subfertile (Blokhina et al., 2021). This makes a sharp contrast with the mouse mutants, which showed complete sterility in males and age-dependent subfertility in females (Herrán et al., 2011). Considering these differences in mutant phenotypes and the temporal expression patterns, teleost *rad21l* might have a function distinct from those in mammals. Interestingly, our experiments in medaka revealed that both *rad21* and *rad21l* transcripts were enriched in the cytosolic fraction of developing oocytes, suggesting the possibilities of their maternal deposition and functions in early embryogenesis (Fig. 2). In zebrafish, scRNA-seq data revealed the same trends as medaka, where *rad21a* and *rad21l* were expressed continuously in a subset of early oocytes (Fig. 3). Notably, zebrafish *rad21l / tp53* double mutants exhibited defects only in the early development but not in the oogenesis itself (Blokhina et al., 2021). In general, early development of non-mammalian vertebrates depends on maternal materials stored in eggs, more than mammals. Further analyses in non-mammalian vertebrates might unveil novel but ancestral functions of these proteins in gametogenesis.

Most of teleost lineages possessed duplicated copies of *rec8* and *rad21* (Fig. 1), while the majority of TGD-derived paralogs have been lost right after their duplication (Inoue et al., 2015). This can be interpreted as follows; both copies of these α-kleisin genes became essential due to functional divergence following their duplication. Concordantly, medaka *rec8a* was expressed in a female-biased manner and its mutants exhibited female-specific sterility (Kikuchi et al., 2019, 2020). A similar pattern was also evidenced in the Nile tilapia, in which *rec8a* had the earlier expression in females, while *rec8b* had the earlier expression and the nuclear localization in males (Luo et al., 2021). Although both were expressed in both ovaries and testes, *rec8a* had cytosolic localization in testes despite its nuclear function, and *rec8b* exhibited no expression in meiotic prophase I oocytes at 30-40 dph, suggesting their sex-specific functions in meiotic prophase I. In this study, we found that the expression bias in *rec8* paralogs was conserved between medaka and zebrafish, which further aligned with the observation in Nile tilapia. In all the three species, *rec8a* and *rec8b* consistently exhibited female- and male-biased expression patterns, respectively, with minor differences in details.

This conserved expression pattern confirms that the bisexual function of the ancestral *rec8* gene was indeed subdivided into each sex in the extant *rec8* paralogs, which likely occurred in the common ancestor of medaka, Nile tilapia, and zebrafish, at least. In addition, there might be the evolutionary constraint preventing those paralogs from reverting to bisexual expression and from losing either of them, for approximately 200 million years. One possible constraint is the stable genomic arrangements of *rec8a* and *rec8b* loci, which ensure the similar expression patterns among teleost species. Another possibility is that the protein function itself diverged into those sexually biased, following the divergence of the expression pattern itself. Future functional evaluation of these paralogs by genome editing may resolve these different evolutionary scenarios.

### Potential origins of sexually biased expression of teleost *rec8* paralogs

Our study revealed the conserved sexually biased expression patterns of *rec8* paralogs (Figs. 2 and 3). Notably, these loci experienced drastic rearrangements after the TGD, and particularly, which likely corresponds to these unique expression patterns (Figs. 1 and 4). We profiled the epigenetic landscape of the mouse *Rec8*, zebrafish *rec8a*, and zebrafish *rec8b* loci and identified putative CREs of these genes (Fig. 4). As a caveat, all the public data used here were obtained from testicular cells rather than ovarian cells, potentially overlooking female-specific CREs. Despite the presence of the testicular CREs, these regions were not highly conserved. Remarkably, those zebrafish CREs have been completely altered at the sequence level even between zebrafish and medaka (Sup. Figs. 3 and 4). This indicates that the conserved expression patterns may originate from lineage-specific CREs, rather than a conserved one. This phenomenon of conserved gene expression despite altered CREs is frequently evidenced in other cases of conserved gene expression, especially those underlying cell type identity (Ogawa et al., 2025; Phan et al., 2025; Sarropoulos et al., 2026). Even with different CREs, a consistent expression pattern is accomplished by TF binding sites (TFBS) turnover, where the same type of TFBS emerges repeatedly at a similar genomic location, presumably due to consistent evolutionary constraints. The sexually biased *rec8* expression might be maintained by TFBS turnover with continuous sexually antagonistic selection, whose relaxed selective pressure might have allowed lineage-specific minor modifications on their expression patterns.

Our TFBS prediction involving chromatin accessibility and motif conservation first highlighted differences in potential regulatory mechanisms between mammalian and teleost *rec8* orthologs (Fig. 5). One of the well-defined gatekeepers of mammalian meiosis is retinoic acid signaling, which activates several different TFs and meiotic genes, via RARs (Koubova et al., 2014; Ishiguro et al., 2020). In agreement with this, RAR binding sites were the most prominent in our analysis for the mouse *Rec8* CREs (Fig. 5). Although those RAR binding sites were found in zebrafish *rec8* CREs, their prediction or conservation scores were not high compared with the other highly-scored TFs (Fig. 5). Germline overexpression of the dominant-negative RAR in zebrafish resulted in the slightly decreased expression level of *rec8a* but not *rec8b*, indicating their weaker but substantial regulatory dependence on retinoic acid signaling compared with mouse *Rec8* (Crespo et al., 2019).

The TFBS analysis also provided insights for the differential regulatory mechanisms between *rec8* paralogs. Overall differences in the highly scored TFBSs between the zebrafish *rec8a* and *rec8b* CREs imply differential *cis*-regulatory codes between paralogs, while this did not pinpoint specific TFs governing their sexually biased gene expression (Fig. 5 and Sup. Fig. 5). FOXL2L is the only TF evidenced to regulate the female-biased expression of the medaka *rec8a* gene (Kikuchi et al., 2019). Our motif analysis targeting zebrafish *rec8* paralogs revealed several potential TFBSs of the Fox-family with relatively high prediction scores at both zebrafish *rec8a* and *rec8b* CREs (Fig. 5). Those Fox-family binding motifs are almost identical to each other, suggesting that any of them have the potential to act as FOXL2L binding sites (Georges et al., 2010; Kikuchi et al., 2019). Of note, *foxl2l* can both upregulate and downregulate target gene expression (Hsu et al., 2025). This implies its repressing function toward *rec8b* in addition to the known function for *rec8a* activation. In line with this, the medaka *rec8b* TSS also harbored Fox-family binding motifs with relatively higher reliability and conservation scores in our preliminary analysis lacking chromatin accessibility data (Sup. Fig. 5). Furthermore, medaka *rec8b* is indeed expressed in XX female individuals under the *foxl2l* knockout background, suggesting the magnitude of the *foxl2l*-*rec8b* regulatory dependency (Sup. Fig. 6). Regarding zebrafish, *foxl2l* is expressed in the same cell types expressing *rec8a* in zebrafish ovaries (Hsu et al., 2025; Liu et al., 2022). Knockout of the zebrafish *foxl2l* gene causes complete female-to-male sex reversal originating from germ-cell dysfunctions and diminished *rec8a* expression, which resembles the phenotype in the corresponding medaka mutants (Nishimura et al., 2015; Ren et al., 2024; Hsu et al., 2025; Yang et al., 2025). This highlights the similar regulatory relationship from *foxl2l* to *rec8a* between medaka and zebrafish despite the absence of conserved CREs. Collectively, the regulatory wiring between *foxl2l* and both *rec8* paralogs might have played a crucial role in the establishment and maintenance of the conserved expression patterns of *rec8* genes.

## Conclusion

In this study, we uncovered the unique α-kleisin gene repertoire in teleosts and the conservation of the sexually biased expression of *rec8* paralogs. These regimes presumably originated from the TGD with its subsequent rearrangements, which might have changed both the evolutionary constraints and the *cis*-regulatory codes of ancestral *rec8* paralogs. These observations highlight the constraints on sexually distinct transcriptional regulation of the essential meiotic genes, which have the potential to disentangle the long-standing mystery of the sex-specific recombination landscape.

## Materials and methods

### Phylogenetic analysis

For phylogenetic tree constructions, peptide sequences of human α-kleisin genes were queried for the NCBI online blastp search against the vertebrate Refseq protein database. Collected peptide sequences were first screened by completeness of their sequences and aligned by the MAFFT online service with default parameters (version 7; Katoh et al., 2019). To maximize the number of phylogenetically informative sites in the alignments, data from cyclostomes (lampreys and hagfishes) were excluded from the primary analysis due to large phylogenetic distances among them. For the same reason, *Rec8* and *Rad21* homologs were separately analyzed first, followed by the analysis involving all chordate *Rad21* and *Rad21l* homologs at once. Unaligned sites were trimmed by trimAl with the automated1 option (v1.4.rev15; Capella-Gutiérrez et al., 2009), and phylogenetic trees were constructed by IQ-TREE (v2.2.6; Minh et al., 2020) with automated model selection. Reproducibility of tree topology was tested by 1000 times bootstrapping tests.

Protein coding genes around each α-kleisin locus were selected for the synteny analysis. Orthologies of these genes were assessed based on the functional annotations by NCBI (National Center for Biotechnology Information) or Ensembl Genome databases. Gene symbols used in this analysis were manually modified (e.g. *fam110a* to *fam110aa* and *fam110ab* in Fig.1e) for the consistency among different strategies for gene symbol nomenclature.

### Animal sampling and sex identification

All animal experiments were conducted under the approval of the Nagoya University official ethics committee (Approval Number 8 and 9 in Department of Science, Nagoya University). The OKcab strain of Japanese medaka was used for the following experiments. *foxl2l* mutant line used in this study was identical to those used in the original study (Nishimura et al., 2015). These fish were maintained in small fresh water tanks at 25 ± 1°C under the light cycle: 14 hrs light and 10 hrs dark. Genetic sex of each individual was determined by PCR as previously described (Kikuchi et al., 2020), which detected the male determining gene *dmy*.

### *In situ* hybridization of medaka gonads

Whole-mount *in situ* hybridization was performed as previously described (Nishimura et al., 2018). In brief, whole bodies of 15-day-post-hatching (dph) XX larvae, testes from six-month-old XY adult individuals, and ovaries from XX adult individuals were used. cDNA clones for *rec8b* (clone name: olte25g11) and *rad21* (olea63c03) were purchased from NBRP Medaka (National BioResource Project Medaka; http://www.shigen.nig.ac.jp/medaka/). *rec8a* and *rad21l* cDNAs were prepared from the total RNA extracted from *meioC* mutant ovaries (Nishimura et al., 2018) and wild-type testes, respectively. These cDNA were amplified by PCR using T7 promoter-tagged primers, followed by amplicon purification from electrophoresed gel. DIG-RNA antisense probes were synthesized using DIG RNA labeling mix with T7 RNA polymerase (Roche). Fixed and permeabilized tissues were mixed with the DIG-RNA probe, labeled with the anti-DIG antibody conjugated with an alkaline phosphatase (Roche), followed by the coloring reaction at room temperature using NBT-BCIP as its substrate. The duration for coloring reaction differed among samples (*rec8a* XY: 10 hrs, other XY: 5 hrs, all XX: 10 hrs). 4 µm-thick sections were prepared by Technovit 8100 kit (Heraeus Kulzer), then counterstained with 1% of neutral red (Sigma-Aldrich). Images were taken under the BZ-X710 microscope (Keyence).

### Single-cell RNA sequencing data analysis of zebrafish gonads

The input sequencing reads of previously published scRNA-seq studies were obtained from the NCBI Sequence Read Archive (SRA) and the Genome Sequence Archive of the National Genomics Data Center, China (Liu et al., 2022; Qian et al., 2022). These datasets were obtained under the accession numbers; SRX13441048 and CRA003925, respectively. The reference genome and its gene annotation were obtained from Ensembl at Ensembl release 109 (GRCz11) and converted into Cell Ranger-acceptable format by CellRanger mkref sub-command (v7.1.0, 10X Genomics). The data used in the following analyses was adopted from the one for ovarian germ cells of 40 dph juveniles. These files were processed into cell-count matrices by the CellRanger. Count matrix datasets were processed by the SoupX (v1.6.2; Young and Behjati, 2020) for ambient RNA contamination and normalized by total UMI (unique molecular identifier) counts or by SCTransform (v0.4.1; Hafemeister and Satija, 2019). Cells were clustered based on their transcriptional profiles, which were visualized by the UMAP method through the Seurat library (v5.0.2; Hao et al., 2024). These clusters were manually annotated by the cell-type specific markers used in the original studies (Liu et al., 2022; Qian et al., 2022).

### ChIP-seq and ATAC-seq data analysis

Previously published results of ChIP-seq for two histone modifications (H3K4me3 and H3K27ac) and ATAC-seq were obtained from NCBI SRA as the fastq file format for both mouse and zebrafish testes. These sequencing reads were trimmed and filtered by their quality using fastp (v1.0.1; Chen et al., 2018) with default options, and mapped onto GRCm38 or GRCz11 reference genomes using bwa-mem2 (v2.3; Vasimuddin et al., 2019). Mapping results were converted into coverage tracks by the bamCoverage utility in Deeptools (Ramírez et al., 2016) with the option “--minMappingQuality 10 --binSize 1 --normalizeUsing CPM” and “--extendReads” for pair-end reads. For the previous ChIP-seq results of DMRT1, RARA, and CTCF, coverage track files were directly obtained from the ChIP-Atlas (Oki et al., 2018; Zou et al., 2022; https://chip-atlas.org/). Accession numbers of all these datasets were attached as Sup. Table 1. These coverage tracks were stacked and visualized by pyGenomeTracks (Lopez-Delisle et al., 2021).

### Regional genome alignment and assessment of nucleotide-level conservation

PhyloP scores of *rec8* loci were obtained from the previous result or calculated by ourselves for mouse and zebrafish, respectively. Previously calculated phyloP scores were based on the genomic alignment of 241 mammalian species performed by the Zoonomia consortium (Foley et al., 2023), and obtained as the bigwig record employing the GRCm38 mouse genome sequence as its reference (https://cgl.gi.ucsc.edu/data/cactus/241-mammalian-2020v2-hub/Mus_musculus/phylop.bw). For zebrafish, genomic regions flanking paralogous *rec8* loci were obtained by the amino-acid-to-nucleotide search of flanking genes against 43 actinopterygian fish genomes by using miniprot (Li, 2023) or manual inspection of the NCBI genome browser. These partial genomic sequences were aligned by progressive-cactus (Armstrong et al., 2020) using the phylogenetic tree obtained from the TimeTree database (https://timetree.org/) and scaled by the previous estimation of neutral nucleotide substitution rates of eight fish species (https://hgdownload.soe.ucsc.edu/goldenPath/danRer7/phyloP8way/fish.mod). The hal-format result was converted into maf-format by cactus-hal2maf utility for the following steps. Based on the zebrafish-based alignment, the neutral substitution model was computed for third codon positions of coding genes in these regions via the phyloFit utility and the conservation scores were computed by the phyloP utility in the PHAST package (Pollard et al., 2010; Hubisz et al., 2011). Regions with low alignment depth (< 4 species) were excluded from phyloP score calculation.

### Prediction for putative transcription factor binding sites

Putative transcription factor binding sites were inferred by the fimo utility in the MEME suite (v5.5.9; Grant et al., 2011) using the JASPER 2020 CORE non-redundant vertebrate motif database. These predictions were applied only for the spans of putative CREs defined in this study (gray shadings in Fig. 4). Prediction scores of mouse and zebrafish were weighted by the prior distributions derived from mapping depth of ATAC-seq reads utilizing the create-priors utility (Cuellar-Partida et al., 2012). Conservation scores of putative binding sites were given as sum of phyloP scores of the sites. These putative binding sites were grouped by the public motif classification released as the JASPAR matrix clustering result for the Vertebrata core motif dataset, and only the site with the lowest *P* value was retained among overlapping binding sites sharing a common motif classification.

### CRediT authorship contribution statement

Taiki Niwa: Writing – original draft, review & editing, Conceptualization, Investigation, Funding acquisition. Mariko Kikuchi: Writing – review & editing, Supervision, Funding acquisition. Minoru Tanaka: Writing – review & editing, Supervision, Conceptualization, Funding acquisition.

## Supporting information

Supplemantary table S1 Accession numbers of public datasets

Supplemantary table S2 PCR primers used in the study

## Acknowledgments

We are grateful to the members of our group of Reproductive Biology in Nagoya University, Mr. Shiyu Kameyama in particular, for their invaluable discussions and supports on animal care, and Dr. Shigehiro Kuraku in National Institute of Genetics for providing computational resources. Computational processes were also performed on the supercomputer system in National Institute of Genetics. We also thank NBRP Medaka for providing cDNAs.

## Data availability

Computational codes and configuration files for genome alignments are available on the github repository (https://github.com/TkNiw/Teleost_aKleisin_codes).

## Supplementary Figures

**Supplementary Figure 1.**
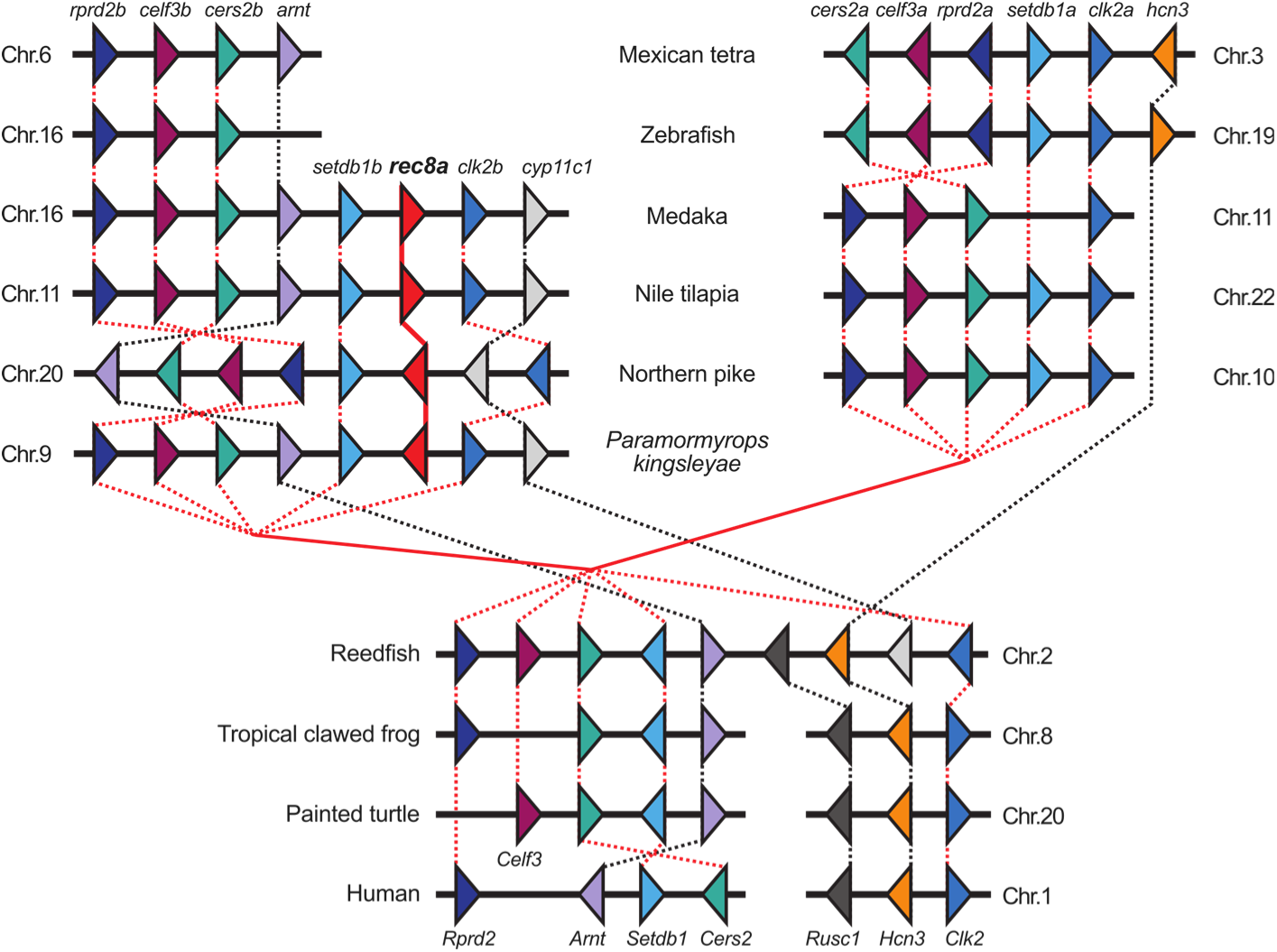
Conserved synteny among *setdb1* loci. Schematic diagrams showing colinear syntenic arrangements of *setdb1* loci, one of which harbors *rec8a* in major teleost lineages. Genes and their transcriptional directions are represented as triangles, and their homology was depicted as edges. Adjacency of genes in this diagram shows their relative positions on the genomes, rather than the actual side-by-side arrangements.

**Supplementary Figure 2.**
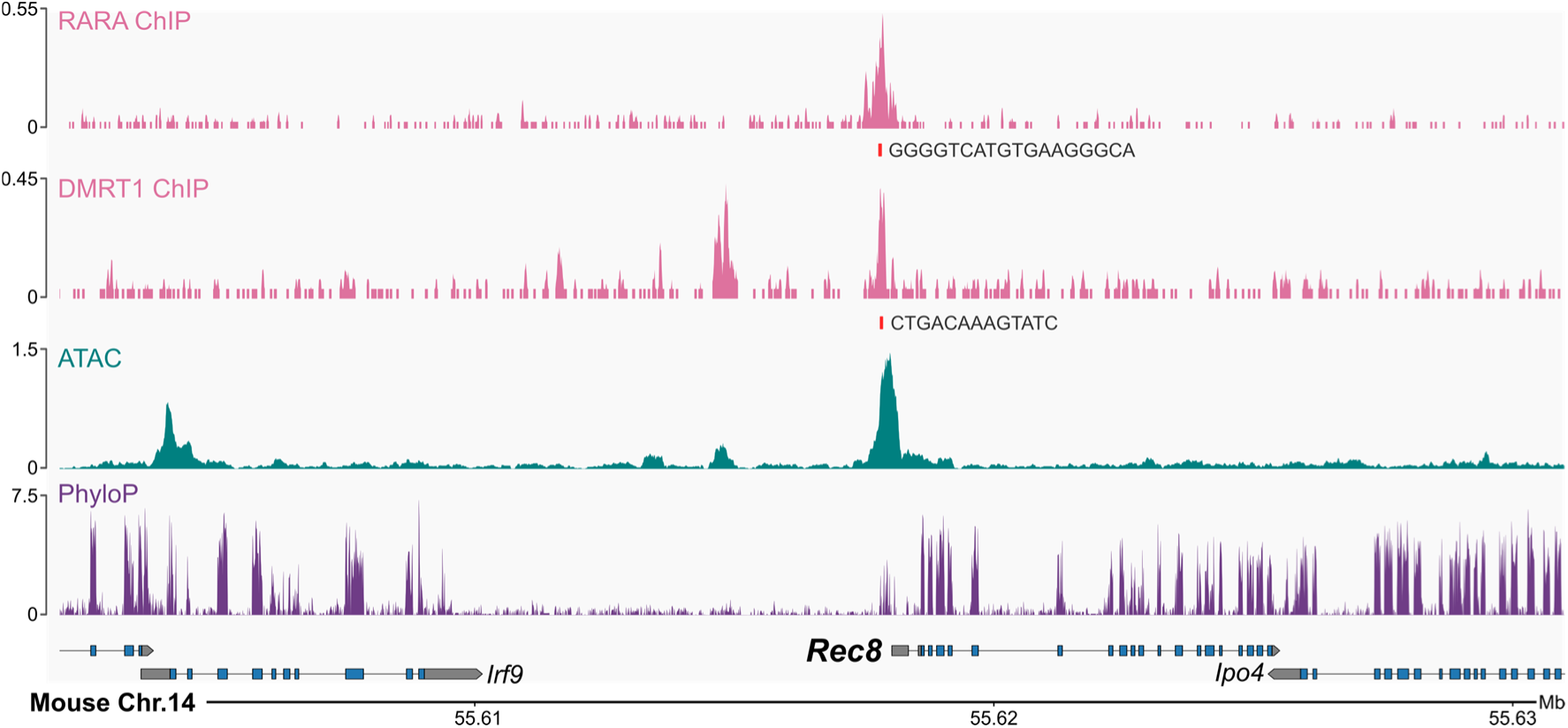
DNA binding profiles of the transcription factors expressed in mouse germ cells. Profiles of DNA binding of two transcription factors in the mouse germline; retinoic acid receptor (RARα) and DMRT1. Read counts were normalized as CPM. Putative binding sites of these proteins were obtained by the result of the prediction tool FIMO with q-value < 0.1.

**Supplementary Figure 3.**
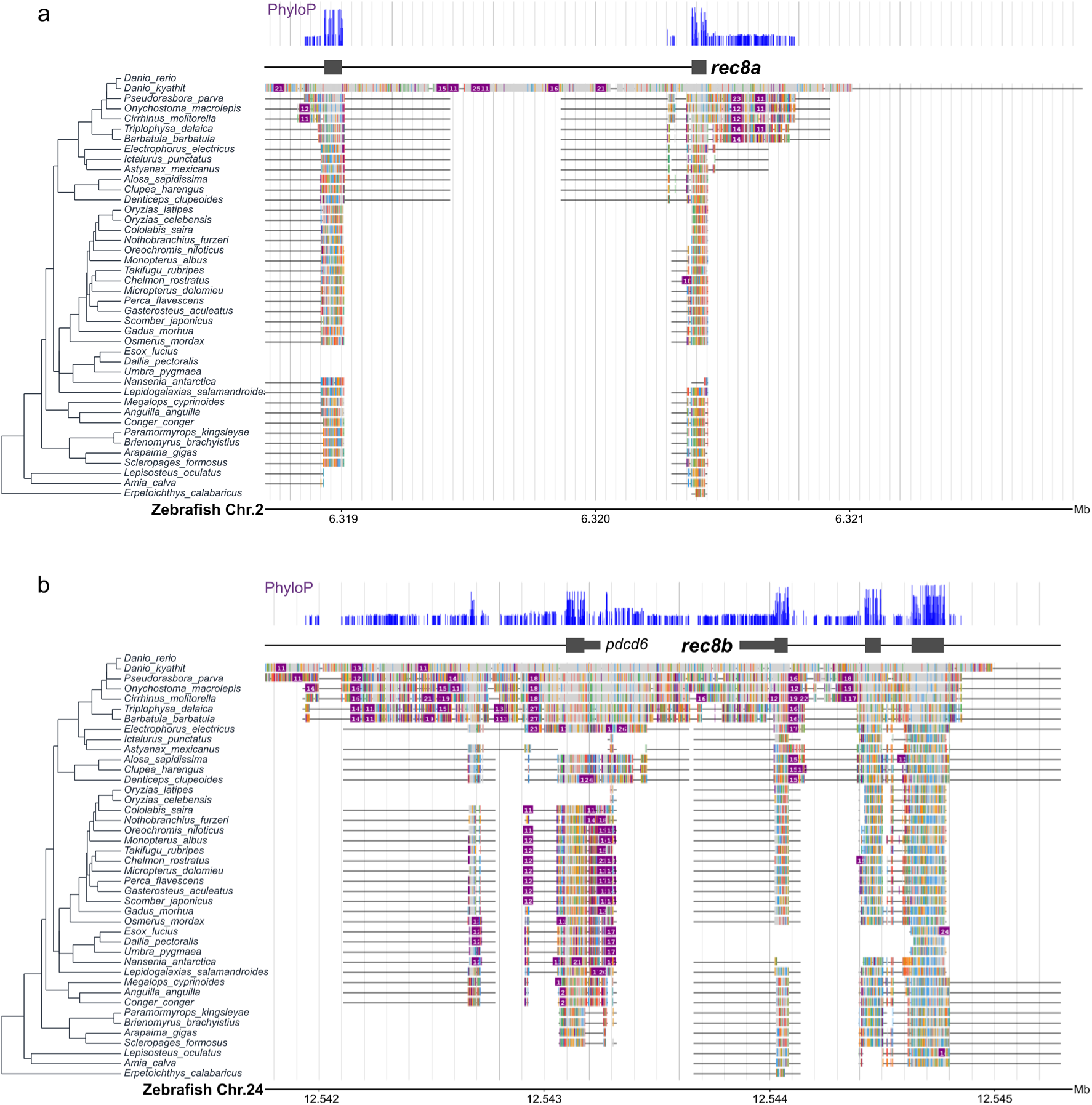
Interspecific genome alignment with the zebrafish reference. Magnified views of the regional genomic alignments around the *rec8a* TSS (a) and the *rec8b* TSS (b). Aligned blocks were indicated as boxes, with different colors for bases unmatching to the reference. Phylogenetic trees are same as the one used for the genomic alignment guide.

**Supplementary Figure 4.**
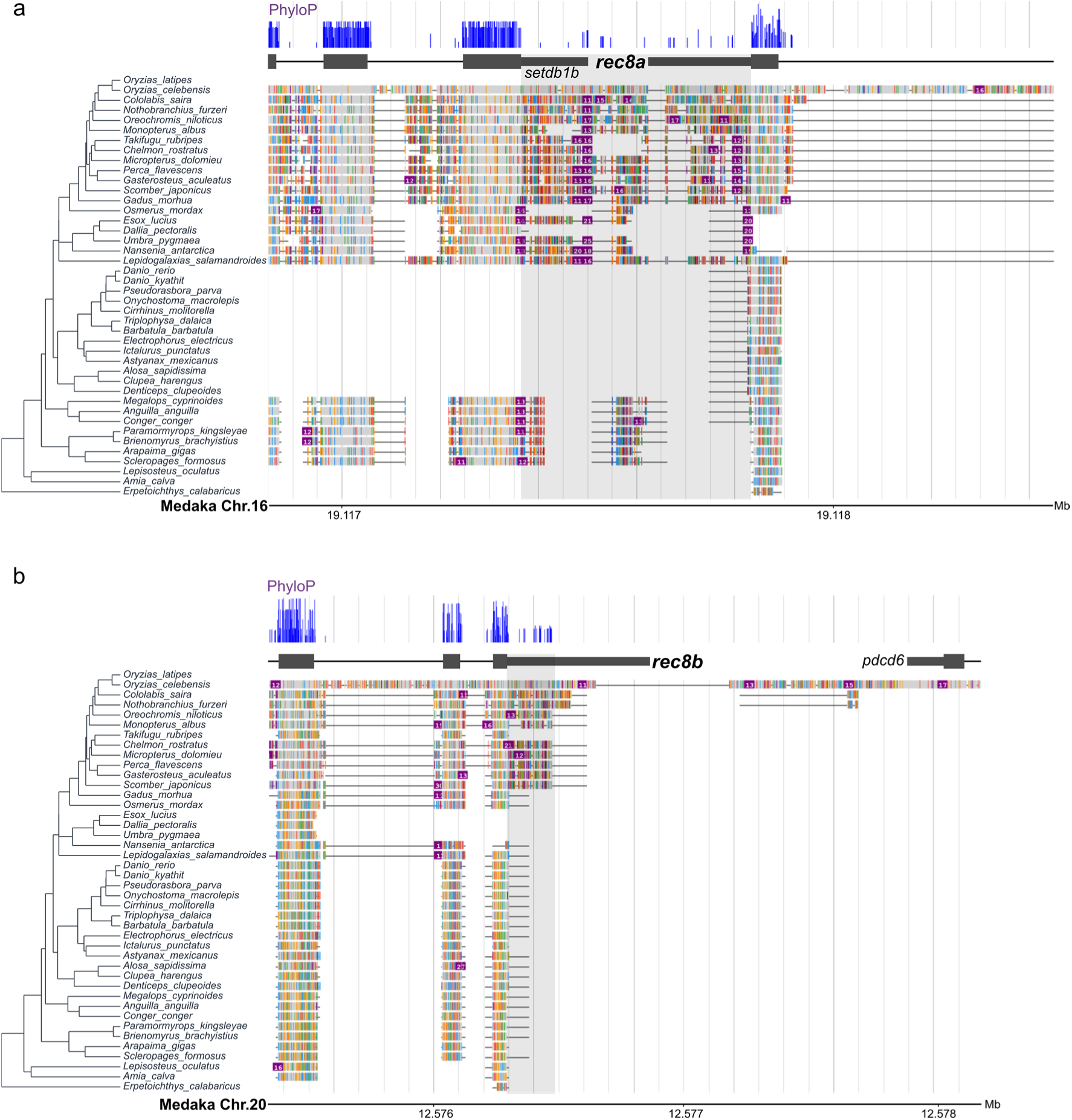
Interspecific genome alignment with the medaka reference. Magnified views of the regional genomic alignments around the *rec8a* TSS (a) and the *rec8b* TSS (b). Aligned blocks were indicated as boxes, with different colors for bases unmatching to the reference. Phylogenetic trees are same as the one used for the genomic alignment guide. Grey shadings show the regions used for the TFBS prediction in Supplementary Figure 5.

**Supplementary Figure 5.**
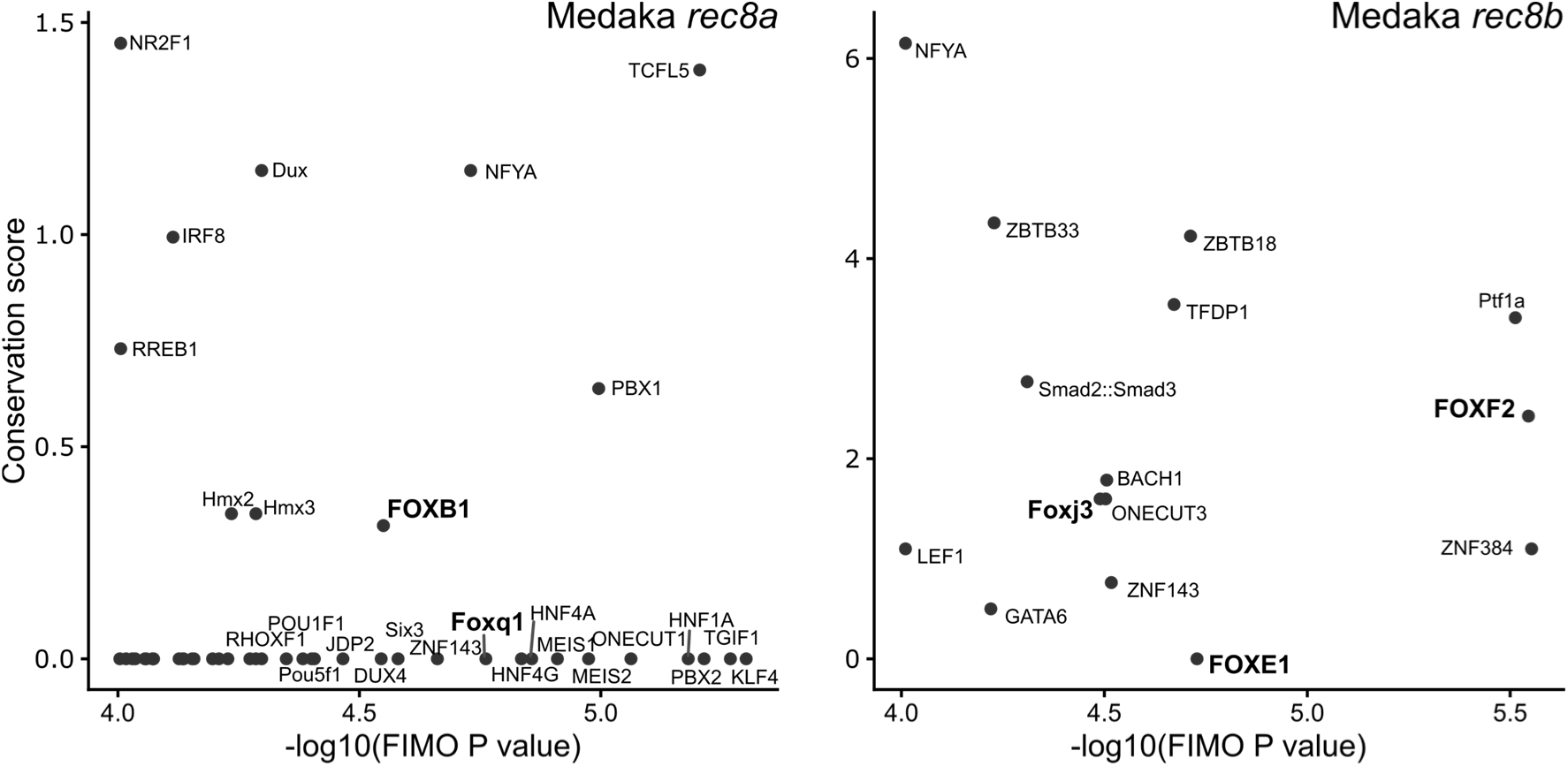
TFBSs inferred within the genomic regions around medaka *rec8a*/*b* TSSs. Horizontal axes show results of TFBS prediction by FIMO only with nucleotide sequences. Genomic regions used for the predictions are indicated in Supplementary Figure 4 as grey shadings in the plots. Conservation score represents the sum of phyloP score at a given motif span. TF names were displayed only for those with the top 20 prediction scores or non-zero conservation scores. TFBSs of the FOX-family were denoted as bold letters.

**Supplementary Figure 6.**
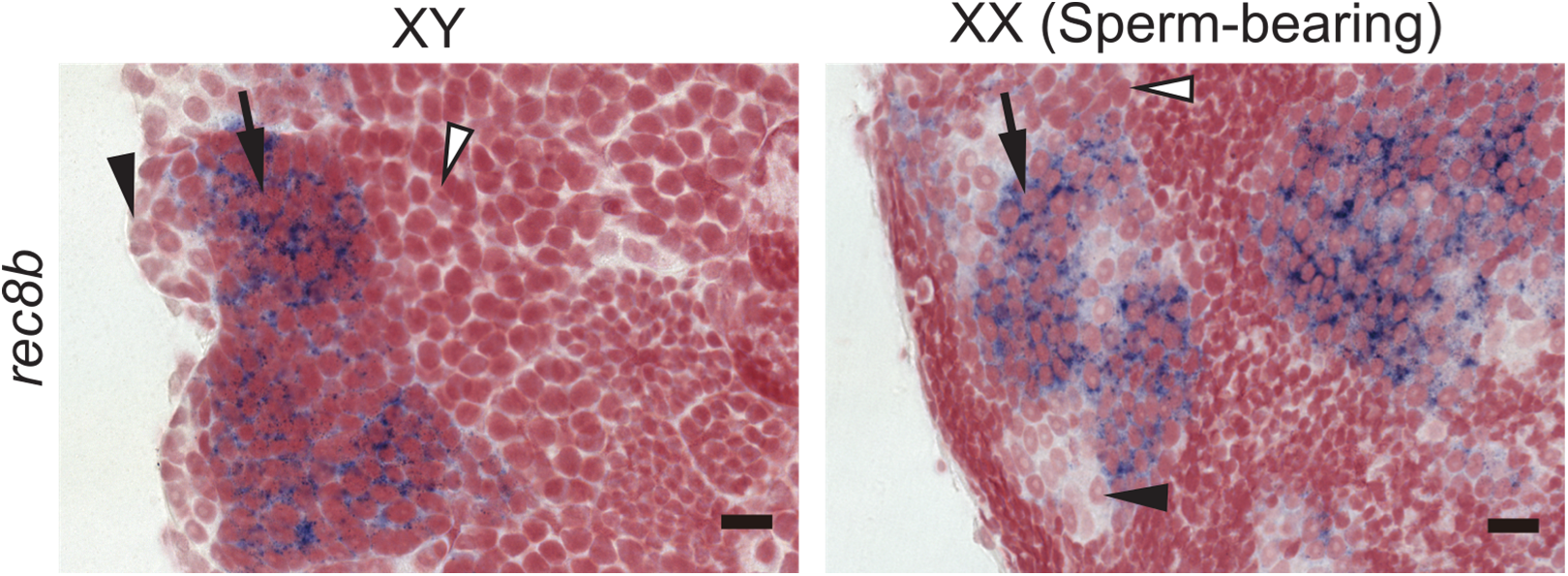
Gonadal expression of *rec8b* in XX female medaka under the *foxl2l* knockout background. Localizations of *rec8b* transcripts on cross sections of adult gonads in XX female and XY male medaka individuals with homozygous *foxl2l*^Δ17^ alleles (Nishimura et al., 2015). Of note, *foxl2l*^Δ17/Δ17^ individuals exhibited complete female-to-male sex reversal of germ cells in XX individuals, while their somatic lineage kept female identity as shown in the original paper. Three individuals were used for each condition; results were consistent among replicates. Arrows and arrowheads show cells with and without signals, respectively. The black and white color of an arrow / arrowheads shows different cell types; mitotic and meiotic cells, respectively. Scale bar: 10 μm.

